# Effects of stimulus pulse rate on somatosensory adaptation in the human cortex

**DOI:** 10.1101/2021.12.04.471210

**Authors:** Christopher L Hughes, Sharlene N Flesher, Robert A Gaunt

## Abstract

**Background:** Intracortical microstimulation (ICMS) of the somatosensory cortex can restore sensation to people with neurological diseases. However, many aspects of ICMS are poorly understood, including the effect of continuous stimulation on percept intensity over time.

**Objective:** Here, we evaluate how tactile percepts, evoked by ICMS in the somatosensory cortex of a human participant adapt over time.

**Methods:** We delivered continuous and intermittent ICMS to the somatosensory cortex and assessed the reported intensity of tactile percepts over time in a human participant. Experiments were conducted across approximately one year and linear mixed effects models were used to assess significance.

**Results:** Continuous stimulation at high frequencies led to rapid decreases in intensity, while low frequency stimulation maintained percept intensity for longer periods. Burst-modulated stimulation extended the time before the intensity began to decrease, but all protocols ultimately resulted in complete sensation loss within one minute. Intermittent stimulation paradigms with several seconds between stimulus trains also led to decreases in intensity on many electrodes, but never resulted in extinction of sensation after over three minutes of stimulation. Additionally, longer breaks between each pulse train resulted in some recovery of the stimulus-evoked percepts. For several electrodes, intermittent stimulation had almost no effect on the perceived intensity.

**Conclusions:** Intermittent ICMS paradigms were more effective at maintaining percepts, and given that transient activity in the somatosensory cortex dominates the response to object contact, this stimulation method may mimic natural cortical activity and improve the perception of stimulation over time.

## Introduction

Intracortical microstimulation (ICMS) of the somatosensory cortex can elicit tactile percepts, even many years after spinal cord injury [1–3]. This can be useful for restoring sensation to people with neurological disease, particularly in the context of a bidirectional brain-computer interface (BCI) [4,5]. The sensations evoked by ICMS can improve robotic arm control by decreasing the time it takes for a person to successfully grasp objects [6]. Apart from functional improvements, ICMS can evoke detectable percepts over many years and stimulation itself does not appear to cause damage that affects neural recordings or detection thresholds [7]. While these factors address the long-term stability and functionality of ICMS, other phenomena may be relevant over short time scales.

One potential issue for sensory feedback via ICMS is percept adaptation. We use the term adaptation to mean a reduction in percept intensity that occurs over time. This effect has been documented for ICMS in the visual cortex [8], stimulation of peripheral nerves [9–11], as well as cutaneous stimulation for sensory substitution [12–14]. Continuous stimulation of peripheral nerves increased the amount of charge required to evoke detectable sensations over time [9–11] and this effect occurred more rapidly at higher stimulation frequencies [10]. However, intermittent stimulation paradigms reduced the effects of adaptation for cutaneous stimulation [13] and in the visual cortex, longer breaks between successive stimulation trains improved the recovery rate of ICMS-evoked percepts [8].

In order for ICMS in the somatosensory cortex to provide a meaningful benefit for people, stimulation will need to provide reliable feedback. Here, we studied the ICMS-evoked perceptual adaptation in the somatosensory cortex to understand the effects of stimulation parameter choices on perceived intensity with the goal of designing better encoding algorithms for bidirectional BCIs.

## Materials and Methods

### Regulatory and participant information

This study was conducted under an Investigational Device Exemption from the U.S. Food and Drug administration, approved by the Institutional Review Boards at the University of Pittsburgh (Pittsburgh, PA) and the Space and Naval Warfare Systems Center Pacific (San Diego, CA), and registered at ClinicalTrials.gov (NCT0189-4802). Informed consent was obtained before any study procedures were conducted. The purpose of this trial is to collect preliminary safety information and demonstrate that intracortical electrode arrays can be used by people with tetraplegia to both control external devices and generate tactile percepts from the paralyzed limbs. This manuscript presents the analysis of data that were collected during the participant’s involvement in the trial, but does not report clinical trial outcomes.

A single 28-year-old male subject with a C5 motor/C6 sensory ASIA B spinal cord injury was implanted with two microelectrode arrays (Blackrock Microsystems, Salt Lake City, UT) in the somatosensory cortex. Data from this participant have been reported previously, including implantation details and initial perceptual effects of ICMS [1], the long-term stability of these devices [7], the effect of ICMS parameters on perception [15,16], and how including ICMS can improve robotic arm control [6]. Each electrode array consisted of 32 wired electrodes arranged on a 6 × 10 grid with a 400 μm interelectrode spacing, resulting in a device with an overall footprint of 2.4 × 4 mm. The remaining 28 electrodes were not wired due to technical constraints related to the total number of electrical contacts on the percutaneous connector. Electrode tips were coated with a sputtered iridium oxide film (SIROF). Additional details on these implants have been published elsewhere [1].

### Stimulation protocol

Stimulation was delivered using a CereStim C96 multichannel microstimulation system (Blackrock Microsystems, Salt Lake City, UT). Pulse trains consisted of cathodal phase first, current-controlled, charge-balanced pulses which were delivered at frequencies from 20-300 Hz and at amplitudes from 2-100 μA. The cathodal phase was 200 μs long, the anodal phase was 400 μs long, and the anodal phase was set to half the amplitude of the cathodal phase. The phases were separated by a 100 μs interphase period. The maximum stimulus train duration for continuous stimulation was 15 s at 100 Hz or 5 s at 300 Hz based on previously defined safety studies [17]. Following any stimulation lasting 15 s, an equal amount of time was spent with no stimulation (50 % duty cycle, 15 s on followed by 15 s off).

### Stimulation adaptation protocols

In the first experiment, the participant used an analog slider to indicate changes in intensity over time while they received continuous stimulation. Analog slide values were scaled from 0-1, representing the minimum and maximum positions of the slider. We considered the intensity to have changed from baseline when the participant moved the slider below 0.95 and the sensation to be completely extinguished when the slider value was less than 0.05. We used these values to account for noise in the analog slider signal. Recordings were taken up to 15 s after stimulation ceased to measure changes in intensity following cessation of stimulation.

We first tested continuous stimulation and delivered pulse trains of 20, 100, and 300 Hz at 60 µA to single electrodes (Fig. 1*A*). We delivered 15 s of stimulation at 20 and 100 Hz and 5 s of stimulation at 300 Hz on 5 electrodes for one trial each.

**Figure 1.**
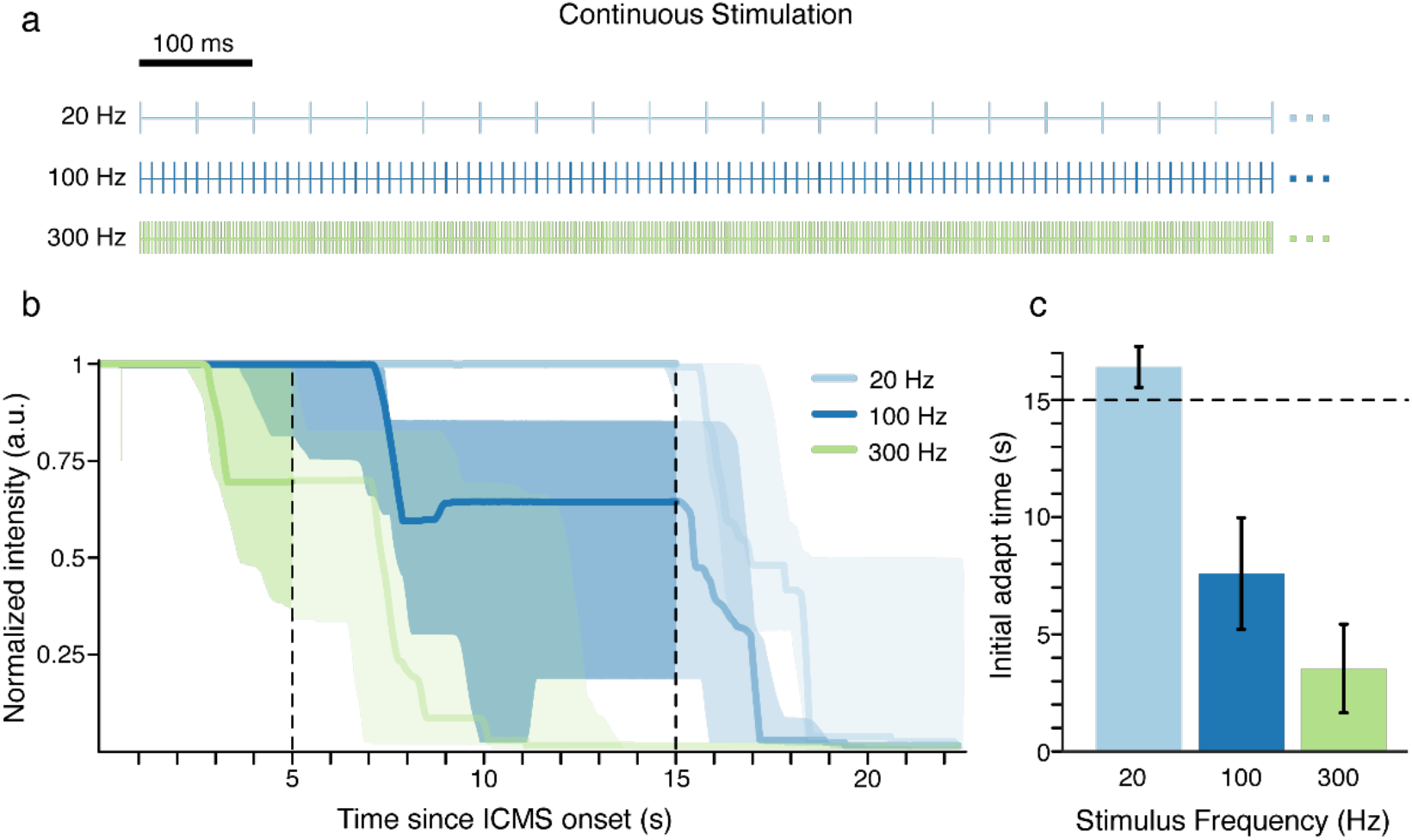
Continuous stimulation at higher frequencies resulted in faster adaptation. A) Continuous frequency trains. Each line represents an individual pulse. B) The participant reported the perceived intensity of the ICMS-evoked sensations using an analog slider. The slider always started at a value of 1 and the participant moved the slider to indicate changes in perceived intensity. Each colored line represents the median intensity for 5 electrodes at a given frequency. The shaded regions represent the interquartile range. The vertical dotted lines indicate the end of stimulation for the 300 Hz train at 5 s and the end of stimulation for the 20 and 100 Hz trains at 15 s. Slider values after stimulation stopped are shown in a lighter shade to emphasize the effects during stimulation. C) Time at which the perceived intensity began to decrease for the median response at each stimulus frequency. Error bars show the estimated standard error and the dotted line indicates the maximum stimulation time of 15 s.

For burst-modulated stimulation, we delivered pulse trains at 100 Hz and 60 µA to single electrodes. To maintain stimulation over longer periods of time, we used a burst modulation scheme with three different burst modulation paradigms: 100 ms, 200 ms, and 500 ms (Fig. 2*A*). Each of these burst modulation protocols consisted of a burst of stimulation followed by a period of no stimulation of equal length. We delivered 60 s of stimulation for each of these burst modulations on 10 different electrodes for one trial each.

**Figure 2.**
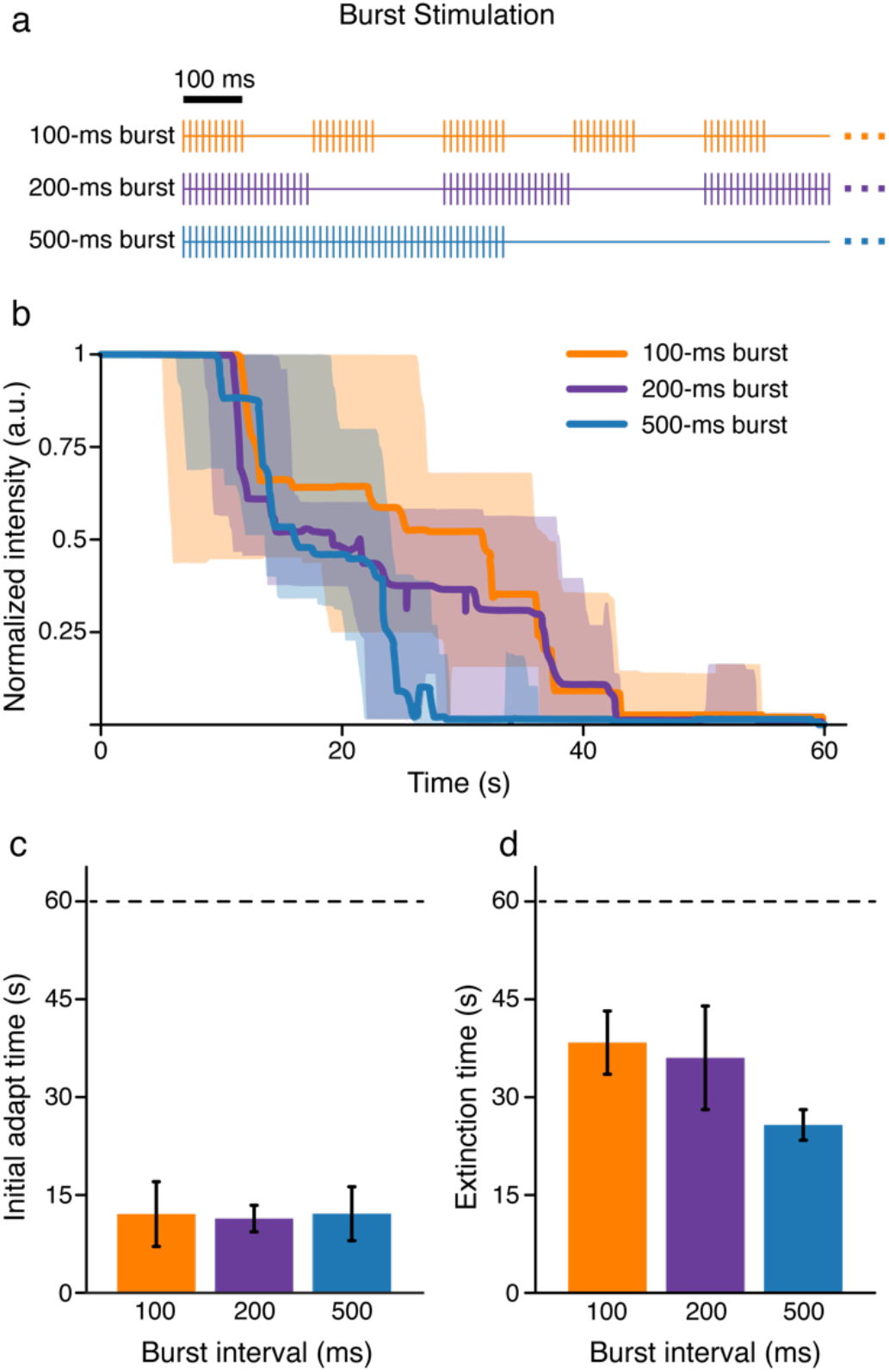
Burst-modulated stimulation extinguished all percepts within 60 s. A) Burst modulated trains. Each line represents an individual pulse. B) The participant indicated the perceived intensity of ICMS with an analog slider. The slider always started at a value of 1 and the participant moved the slider to indicate changes in perceived intensity. Each line represents the median intensity value across 10 tested electrodes for a given burst length. The shaded regions represent the IQR. C-D) Bar plots showing C) the time at which the percept intensity began to decrease and D) the time at which the percept became was extinguished. Error bars show the estimated standard error. The dotted line indicates the maximum stimulation time of 60 s.

For intermittent stimulation trials, there was an initial adaptation period followed by a recovery period. In the adaptation period, the participant received 1 s of stimulation at 60 µA followed by 5 s of no stimulation for 50 repetitions to measure changes in intensity over time (Fig. 3*A*). After each 1 s pulse train, the participant verbally reported the perceived intensity on a self-selected scale. The participant had performed similar magnitude estimation tasks in previous studies, and used consistent ratings for the same parameters and electrodes across sessions [15]. We had the participant use a self-selected scale rather than normalizing to the first value because we were interested in whether the magnitude of the initial perceived intensity on different electrodes related to the change in intensity over time. After the adaptation period, the participant received 1 s of stimulation followed by 61 s without stimulation for another 5 repetitions. We used this period to assess any recovery in the perceived intensity over time.

**Figure 3.**
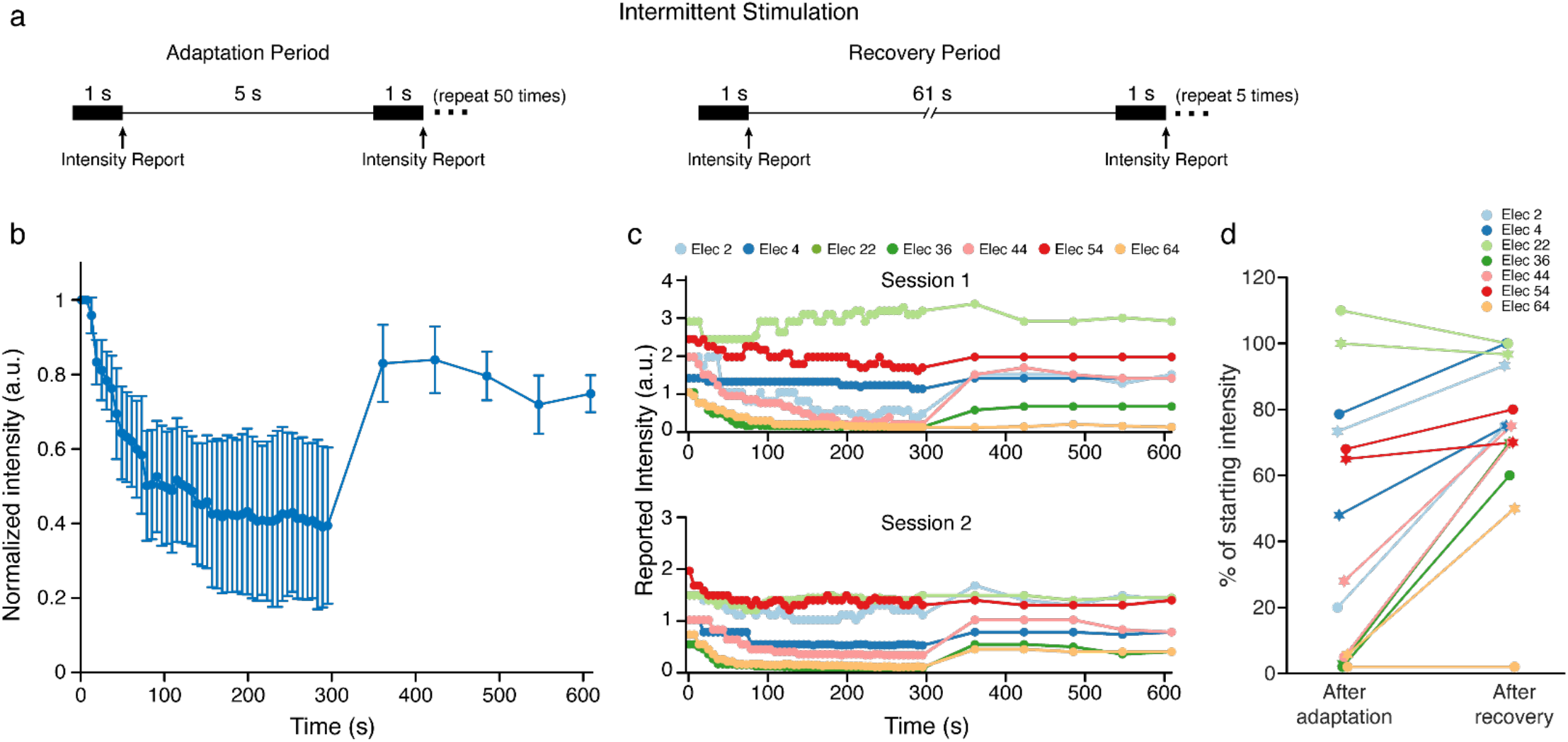
Intermittent stimulation caused less adaptation, which partially recovered over time. A) Intermittent adaptation and recovery paradigms. The participant indicated the perceived intensity for each stimulus train directly following each stimulation train. B) Blue dots represent the median intensity value measured at each time point across all electrodes. Error bars show the estimated standard error. (C) The response of each electrode in two sessions during the adaptation and recovery periods. (D) The percent change in intensity for each electrode after the adaptation and recovery periods. The change in intensity is calculated from each electrode’s initial intensity. Circular markers indicate the first session while star markers indicate the second session.

Seven electrodes were tested twice with the intermittent paradigm at 100 Hz to measure differences in adaptation over time between electrodes. Four electrodes were also tested four times with the intermittent paradigm at 20, 100, and 250 Hz to measure differences in adaptation as a function of stimulation frequency.

### Detection threshold adaptation protocol

We measured detection thresholds using a two-alternative forced choice task. Two intervals were presented with one containing stimulation. The participant was asked to report which interval contained the stimulus. Amplitude was modulated using a one-up three-down staircase method [18,19]. The initial current amplitude was 10 μA and was increased or decreased by a factor of 2 dB with a constant frequency of 100 Hz. After five changes in the direction of the current amplitude (increasing to decreasing, or decreasing to increasing), the trial was stopped. The detection threshold was calculated as the average of the last 10 values tested before the fifth direction change. Following the first detection task, the participant received stimulation using a 15 s on 15 s off protocol – the maximum time we can stimulate continuously – at 60 µA and 100 Hz for 240 s. The detection task was then repeated. We conducted this task on 12 different electrodes each with one trial.

### Statistics

All statistical analysis was conducted in MATLAB (Mathworks, Natick, MA). We used linear mixed effects models to test for relationships among the variables of interest (fixed effects) while excluding the impact of variables not of interest (random effects).

For the continuous and burst stimulation paradigms, we defined the time at which the intensity fell below 0.95 to be the initial adaptation time. To understand how intensity decreased as a function of frequency for the intermittent stimulation paradigm, we compared the intensity after the first stimulus train of the adaptation period to the intensity after the last stimulus train of the adaptation period. We repeated this analysis for the recovery period by comparing the intensity of the last repetition of the adaptation period to the last repetition of the recovery period.

To determine if the ability to detect stimulation changed with long periods of stimulation, we compared the detection thresholds measured before and after a long period of continuous stimulation. Reported p-values are for the coefficient of the fixed effect and were considered to be significant at α = 0.05.

The analog slider values are plotted as the median and the interquartile ranges (IQR). Data from other experiments were also found to be non-normal using the Anderson-Darling test. For all other analysis we used bootstrapping to calculate the median from a random sampling of each data set with replacement and repeated this 10000 times, resulting in 10000 estimated medians. The mean and standard deviation of these bootstrapped samples were used to estimate the population median and standard error [20].

## Results

### Continuous stimulation at higher frequencies results in faster adaptation

We delivered continuous stimulation for 15 seconds at 20 and 100 Hz and for 5 seconds at 300 Hz on 5 electrodes (Fig. 1*A*). The time at which the intensity began to decrease was significantly different between different stimulus frequencies (p = 8.1e-5, Fig. 1*B,C*). 300 Hz stimulation resulted in the fastest change from baseline, with the median intensity falling to 69% of the baseline intensity after 5 s of stimulation. The median intensity remained unchanged from baseline after 5 s of stimulation at both 20 Hz and 100 Hz. However, starting 7 s after stimulation onset, 100 Hz stimulation caused changes in percept intensity that fell to 64% after 15 s of stimulation. Stimulation at 20 Hz had no effect on intensity during the 15 s stimulation window. Ultimately, higher stimulation frequencies caused faster adaptation (Fig. 1*C*).

### Burst stimulation extinguishes sensation over short periods

To potentially extend the time ICMS could be provided, we tested burst-modulated ICMS using 100 Hz pulse trains for 60 seconds. Burst modulation stimulus trains with a 50% duty cycle and burst durations of 100 ms, 200 ms, and 500 ms were tested (Fig. 2*A*). All burst modulation schemes delivered the same number of pulses in 60 s. Stimulation for 60 s caused a complete extinction of the evoked sensation on 26 of the 30 trials (Fig. 2*B*) and there was no difference in the time at which the perceived intensity began to decrease between burst paradigms (p = 0.76, Fig. 2*C*). However, there was a difference in the time required for the sensations to become undetectable between the three burst stimulation paradigms, with the 500 ms paradigm causing the fastest extinction (p = 0.012, Fig. 2*D*).

Compared to continuous stimulation at 100 Hz, the initial adaptation during burst stimulation happened more slowly, but this difference was not significant (p = 0.13). Ultimately, decreasing the burst duration extended the useful perceptual window, but the intensity always decreased over time, and regardless of the burst duration, sensations were completely extinguished over relatively short periods of time (Fig. *2B,D*).

### Increasing time between stimulus trains preserves evoked percepts

Next, we tested an intermittent stimulation protocol with larger gaps between stimulation trains. This protocol was divided into an adaptation period and a recovery period. In both periods the stimulus trains were 1 s long, but the gap between trains was increased from 5 s in the adaptation period to 61 s in the recovery period (Fig. 3A) to allow us to measure whether percept intensity recovered. With this protocol, the median percept intensity still decreased during the adaptation period, however the percepts were not completely extinguished and also increased in intensity during the recovery period (Fig. 3*B*). Additionally, while the percepts became very weak on some electrodes during the adaptation period, the participant still reported feeling them and no electrodes ever became imperceptible (Fig. 3*C*). Considering all of the electrodes together, the median intensity decreased to 40% of the initial intensity during the adaptation period and recovered back to 75% of the initial intensity by the end of the recovery period (Fig. 3*B*).

There was considerable variability between the electrodes in how much the percept intensity adapted and recovered (Fig. 3*C*). In both sessions, the sensations on two electrodes were nearly extinguished – the perceived intensity decreased to less than 6% of the initial intensity – during the adaptation period (Fig. 3*C,D*, Elec 36 and Elec 64) with varying amounts of recovery. On the other hand, the intensity on two electrodes decreased to 48-79% (Fig. 3*C,D*, Elec 4 and 54), while one electrode showed no change in intensity (Fig. 3*C,D*, Elec 22).

### Decreases in intensity of intermittent stimulation are consistent across frequencies

We repeated the intermittent stimulation paradigm on four electrodes at 20 Hz, 100 Hz, and 250 Hz (Fig. 4). These three frequencies represent low, intermediate, and high frequencies within our stimulus range. Consistent with previous findings [15], we found that frequency had electrode specific effects on the evoked intensity; stimulation on one electrode elicited the highest intensity at the lowest frequency (Fig. 4A, Elec 54) while stimulation on the other three electrodes elicited the highest intensity at the highest frequency (Fig. 4A, Elec 4, 22 and 36), resulting in different initial intensities at different frequencies. The effect of frequency on intensity changes during the adaptation period were minimal in this latter group of electrodes. However, on electrode 54, stimulation at higher frequencies led to more adaptation (Fig. 4*B*), although the intensity decreases during the adaptation period and increases during the recovery period were not significantly different across frequencies for any electrode (p >= 0.05, Fig. 4*B*). This indicates that the magnitude of the change in intensity was constant, despite the different initial intensities.

**Figure 4.**
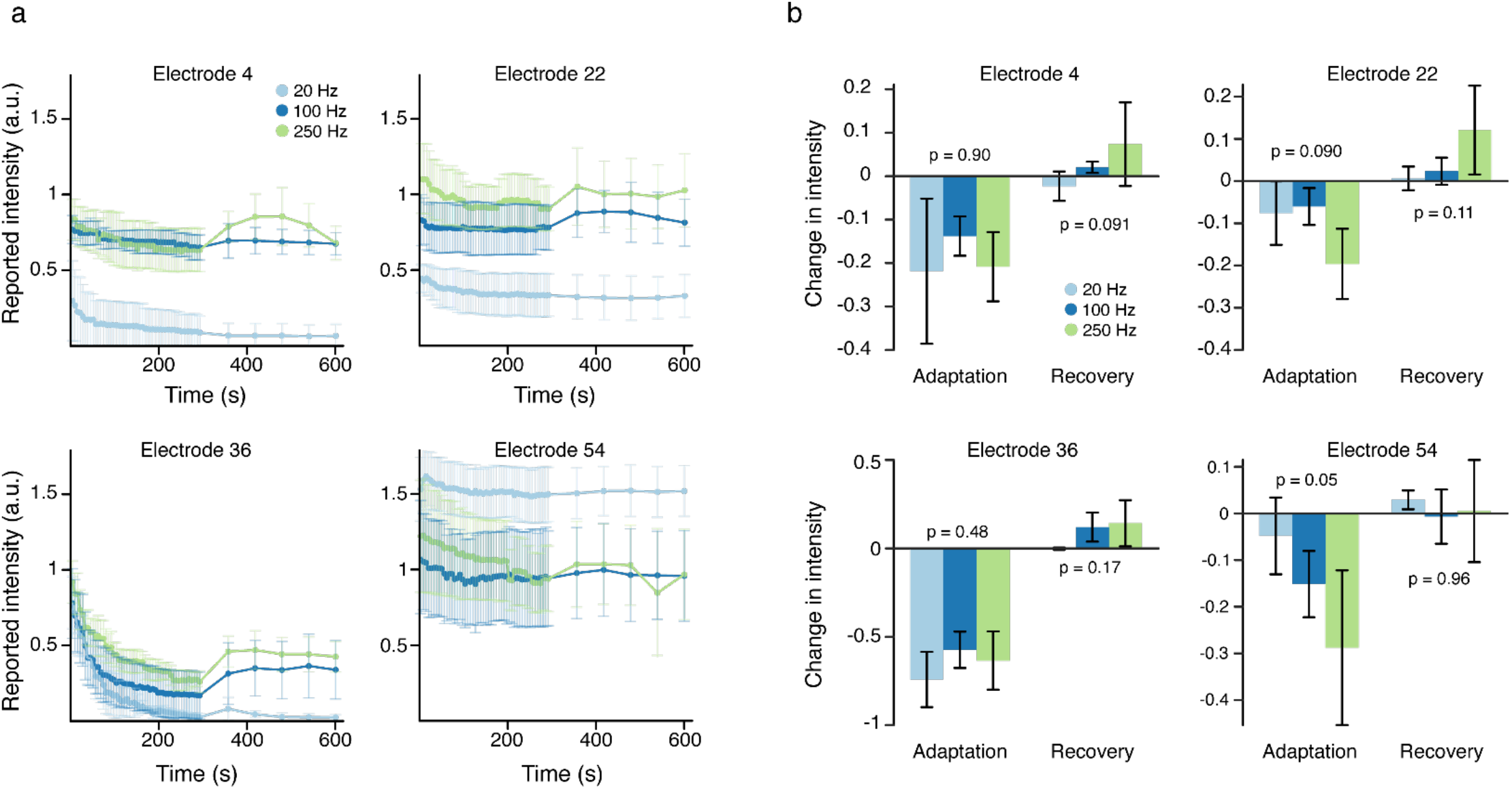
Stimulation frequency did not affect adaptation and recovery on individual electrodes. A) Using the intermittent stimulation protocol, four different electrodes were stimulated at 20, 100, and 250 Hz. Colored dots represent the median intensity value measured at each time point across 4 test sessions. Error bars show the estimated standard error. Different colors indicate the different stimulation frequencies. B) Intensity changes during adaptation and recovery at the three different frequencies. The change in intensity for the adaptation period was the measured between the beginning and end of the adaption period while the change for the recovery period was measured between the between the end of the adaptation period and end of the recovery period. Each bar shows the median difference across four test sessions. Error bars show the estimated standard error.

### Detection thresholds did not charge after continuous stimulation

We also measured the detection threshold before and after long periods of stimulation. We measured the detection threshold and then delivered 4 minutes of stimulation using 15 s of stimulation followed by a 15 s break, which was the maximal length of time we could continuously deliver stimulation. We then remeasured the detection thresholds. Using this protocol, we found no significant difference in the thresholds measured before and immediately after stimulation across 12 electrodes (p = 0.10, Fig. 5) with a median threshold increase of 1.3 µA following continuous stimulation.

**Figure 5.**
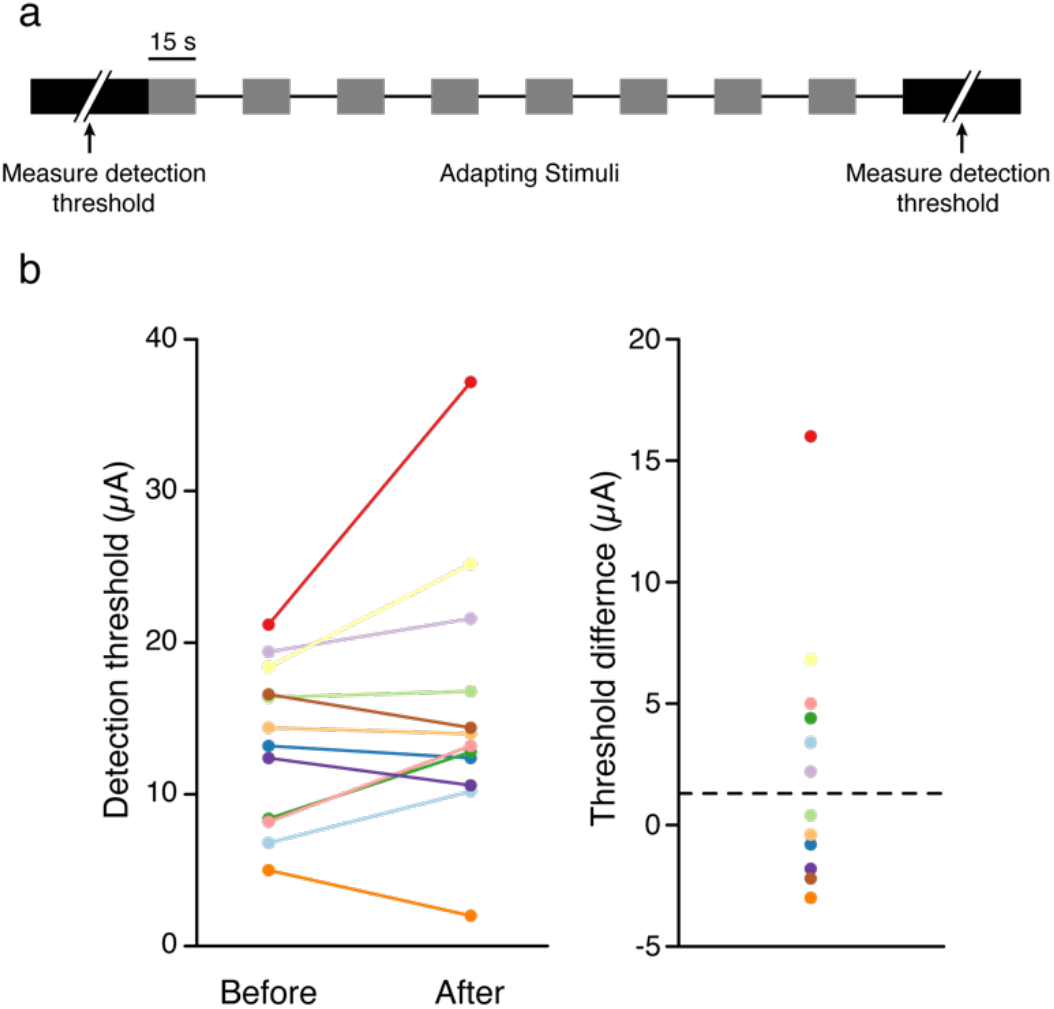
Long periods of stimulation did not have a significant effect on detection thresholds. A) We measured the detection threshold and then applied stimulation for 15 s at 100 Hz followed by 15 s of no stimulation for four minutes and then measured the detection threshold. B) Detection thresholds before and after adaptation protocol. Each color represents a different electrode. C) The difference in detection thresholds before and after the adaptation paradigm. Colors are the same as in panel b. The dotted line represents the median threshold difference.

## Discussion

### Effect of stimulation frequency on adaptation

Continuous stimulation at high frequencies as well as burst-modulated stimulation protocols caused adaptation of the perceived intensity, and eventually extinction of the sensations for burst-modulated paradigms. However, at the lowest stimulation frequency there was no change in the perceived intensity over 15 s of stimulation. This is analogous to observations in the peripheral nervous system in which lower frequencies increased the time before intensity decreases occurred [10], although this has not been true for all studies [9]. More rapid adaptation with higher frequencies has also been noted for electrocutaneous stimulation [12,14] as well as for vibrotactile stimulation [21–25] where higher frequencies resulted in larger and faster changes in detection thresholds as well as decreases in the ability to discriminate stimuli. For electrocutaneous stimulation, stimulus rates over 1000 Hz led to sensation extinction within seconds [14], while lower frequencies and burst paradigms allowed sensations to be evoked for many minutes [12].

Here, the only stimulus paradigm that that did not cause adaptation was continuous stimulation at 20 Hz for 15 s. Due to protocol limitations, driven by concerns about stimulation-driven tissue damage [26,27], we do not stimulate continuously for periods longer than this, so we cannot know if these low frequencies would have caused adaptation over longer periods of time. While this is encouraging, there are limitations using low frequency stimulation. Low frequency stimulation typically evokes qualities of “tapping” or “sparkle” that could be undesirable for object grasping, and on some electrodes low frequencies are unable to drive sensation at all [15]. Therefore, low frequency stimulation itself will not provide a practical means to provide reliable sensations for all electrodes and qualities.

### Intermittent stimulation reduced adaptation

Intermittent stimulation allowed sensations to be evoked over much longer periods of time than continuous stimulation, similar to electrocutaneous stimulation [13]. While adaptation still occurred with intermittent stimulation, the magnitude and duration were electrode dependent, similar to the effects of ICMS in the visual cortex [8]. Interestingly, these electrode-dependent effects ranged from stimulation nearly eliminating the percepts, to having no effect at all (Fig. 3*C*). Importantly, intermittent stimulation never completely extinguished the sensations over 200 s using 100 Hz stimulus trains. Intermittent stimulation conceptually mimics the transient neural activity of many cortical neurons during touch [28] and could provide stimulation during important task-dependent intervals, evoking more reliable sensations over longer periods of time.

### Detection thresholds did not change after long periods of stimulation

Although both continuous and intermittent stimulation affected the perceived intensity, we did not find any significant effect on the detection thresholds. This stands in contrast to results for peripheral nerve and electrocutaneous stimulation, where detection thresholds increased significantly after continuous stimulation [9,12]. Furthermore, this result appears to be inconsistent with the decreases in intensity that occurred. However, there are two relevant factors to consider.

First, detection thresholds and intensity perception are not necessarily measures of the same phenomenon [15]. We found that in the human somatosensory cortex higher stimulation frequencies always decreased detection thresholds, but had variable effects on perceived intensity at suprathreshold amplitudes. Second, measuring detection thresholds took 2-5 minutes and consisted of 1 s stimulus trains followed by a delay of 5-10 seconds for the null stimulus interval and participant response. This made the detection threshold protocol similar to the intermittent stimulation protocol, which itself caused adaptation. In fact, we found that changes in percept intensity driven by intermittent stimulation stabilized after approximately 100 s (Fig. 4). This would mean that the detection task alone could drive adaptation. Unfortunately, any task used to calculate detection thresholds necessarily requires stimulation, making the direct effect of stimulation on thresholds difficult to determine.

### Physiological mechanisms of adaptation

Adaptation of the perceived intensity and changes in the ability to both discriminate and detect tactile stimuli occur in normal touch [21–25,29–33] and are most likely driven by central mechanisms [33]. This adaptation occurs in multiple sensory cortices, where a constant stimulus typically results in rapidly decreasing neural responses [34,35]. Additionally, adaptation of neural responses has been observed for ICMS in mouse cortex, where high-frequency stimulation led to rapid adaptation of neurons away from the electrode [36]. One possible mechanism, short-term depression, has previously been implicated in adaptation, specifically for thalamocortical projections [37]. Short-term depression at thalamocortical synapses was purported to play a strong role in rapid adaptation to brief stimuli provided to the whisker or electrical stimulation (adaptation and recovery within seconds), but separate mechanisms were suggested for slow adaptation (adaptation for minutes or longer). Another possible mechanism is inhibitory neuron drive [38–43]. Two specific types of inhibitory interneurons, parvalbumin and somatostatin neurons, play important roles in adaptation in the auditory cortex [42,43], with parvalbumin neurons providing continuous inhibition throughout the stimulus and somatostatin neurons providing dynamic inhibitory drive based on the number of repetitions.

Taken together, the literature suggests that adaptation occurs as a normal part of cortical processing of sensory information. Slow adaptation may depend more on the activity of different neuronal subtypes while rapid adaptation depends more on short-term depression at synapses. This could explain why continuous ICMS led to consistent decreases in intensity while intermittent ICMS had more electrode specific effects. Electrode specific effects may be related to the density of different neuronal subtypes recruited by ICMS. Because ICMS bypasses subcortical inputs, adaptation that occurs from ICMS may look different than normal sensory adaptation and may depend on the targeted area and stimulated electrode. Normal tactile input leads to changes in intensity, but not extinction of perception. Continuous ICMS leads to extinction, implying that adaptation driven by ICMS is not identical to adaptation driven in normal sensory processing.

### Limitations

This study represents the first study of adaptation to ICMS in human somatosensory cortex. However, there are several limitations that should be considered. First, these experiments were only conducted in one participant. Additional data in other participants will be needed to understand if there are participant or implant location specific effects. Additionally, there is a very large parameter space (electrode, amplitude, frequency, burst intervals, etc.) and only a few specific parameter combinations were considered here. One reason for this is that there is typically limited experimental time available with human participants, making it difficult to explore comprehensive parameter sets. Further, we were not able to change pulse width because of limitations in our protocols. Data collected in more participants with additional parameter variations will provide further insights into the nature of adaptation. Working in humans also limits our ability to directly measure neural activity during stimulation, limiting our ability to comment on neural mechanisms.

### Implications for bidirectional brain-computer interfaces

There are at least two approaches to encoding sensory information in stimulus trains: biomimetic [44–46] and engineered [6,47,48] encoding. Biomimetic approaches aim to create patterns of stimulation that mimic the neural activity that occurs during natural touch, while engineered approaches aim to provide informative stimulation that can be learned to represent specific inputs. Both approaches have met with success in applications in the peripheral nervous system for improving robotic arm control [44,45,48,49]. Biomimetic feedback in the peripheral nervous system evoked more natural sensations and improved performance on some motor tasks [45]. Biomimetic stimulation has not been tested for ICMS, however, a simple encoding in which force was linearly transformed to amplitude improved neuroprosthetic control [6]. However, if contact is maintained for long periods of time with this paradigm, the percepts will become undetectable after just a few seconds (Fig 2). Burst stimulation and lower frequencies could help extend the percept time, but with limitations.

Biomimetic encoding may drive more natural adaptation processes. Biomimetic pulse trains can be built from computational models that predict neural activity based on tactile input [50–52]. These models predict that during object contact, large populations of neurons become active with high firing rates. During maintained contact, the number of active neurons and their firing rates are significantly reduced, which aligns with recorded neural activity in the cortex during touch [28]. Biomimetic stimulus trains would naturally resemble the intermittent stimulation protocols tested here (Fig. 3), allowing sensations to persist over longer periods of time without extinction. We suggest that biomimetic approaches may provide a reliable way to provide sensory feedback for bidirectional BCI applications and may align more directly with normal cortical activation and adaptation.

## Funding

This work was supported by the Defense Advanced Research Projects Agency (DARPA) and Space and Naval Warfare Systems Center Pacific (SSC Pacific) under Contract N66001-16-C4051 and the National Institute of Neurological Disorders and Stroke of the National Institutes of Health under Award Numbers UH3NS107714 and U01NS108922. SNF was supported by an NSF Graduate Research Fellowship under grant number DGE-1247842. Any opinions, findings and conclusions or recommendations expressed here are those of the authors and do not necessarily reflect the views of DARPA, SSC Pacific, or the National Institutes of Health. The funders had no role in the study design, data collection, interpretation of the results, or the decision to submit this work for publication.

## Data Statement

Data and code are available upon reasonable request.

## CRedIT authorship contribution statement

**Christopher L. Hughes:** Conceptualization, Methodology, Software, Formal Analysis, Investigation, Writing-Original Draft, Writing-Review & Editing, Visualization. **Sharlene N. Flesher:** Conceptualization, Methodology, Software, Investigation, Writing-Review & Editing. **Robert A. Gaunt:** Conceptualization, Methodology, Writing-Review & Editing, Visualization, Supervision.

## Declaration of competing interest

None.

## Acknowledgements

We would like to thank N. Copeland for his extraordinary commitment to this study, as well as Debbie Harrington (Physical Medicine and Rehabilitation) for regulatory management of the study.

## Notes

### Competing Interest Statement

The authors have declared no competing interest.

